# Emotional contexts influence vocal individuality in ungulates

**DOI:** 10.1101/2024.09.18.613506

**Authors:** Anna N. Osiecka, Romain Lefèvre, Elodie F. Briefer

**Affiliations:** University of Copenhagen, Behavioural Ecology Group, Section for Ecology and Evolution, Department of Biology, 2100 Copenhagen, Denmark; Austrian Academy of Sciences, Acoustics Research Institute, 1040 Vienna, Austria; University of Vienna, Department of Behavioral and Cognitive Biology, 1030 Vienna, Austria; University of Saint-Etienne, ENES Bioacoustics Research Laboratory, 42000 Saint-Etienne, France

**Keywords:** affective states, contact calls, information coding, information loss, vocal behaviour

## Abstract

To group-living animals, such as most ungulates, being able to recognise the members of one’s social groups is crucial. While vocalisations often carry cues to identity, they are also impacted by the affective state of the caller, with signals often becoming more chaotic in contexts of negative valence or high arousal. How might this influence vocal individuality – and is there a pattern across taxa? To understand how the individual information content is maintained over emotionally charged contexts, we studied putatively negative and positive contact calls of seven ungulate species: cattle, goats, sheep, pigs, wild boars, horses, and Przewalski’s horses. The information content of these calls was assessed using Beecher’s statistic and the Potential of Individuality Coding. Across the species, the information content was lower in calls of negative than positive valence. In each of the species, at least one acoustic parameter was a reliable indicator of individuality across contexts. Our results indicate that negative valence overrides individual information in ungulate vocalisations at least to some extent, and imply that individual vocal recognition may require acoustic stability of certain important parameters. These findings reveal a nuanced role of affective communication in maintaining social bonds among socially complex animals.

## Introduction

In many types of social groups (e.g. family and extended family groups), some form of individual recognition is needed to achieve a good functioning of daily tasks (Ward & Webster 2016). Acoustic cues are often favoured for that purpose since they can travel far and across obstacles, and may be easier to use in a crowd than e.g. olfactory cues (often requiring direct contact; Tibbetts & Dale 2007). Across species, cues to caller identity can be coded in various spectral, temporal, and amplitude features, and carried by both stereotyped and variable vocalisations (see review in Wyman et al. 2022).

Yet vocalisations do not exclusively carry information on identity, but may instead inform on multiple aspects of the sender, such as their sex (e.g. Cordeiro et al. 2012, Volodin et al. 2016), size (e.g. Briefer & McElligott 2011a, Wyman et al. 2012), or physiological state (e.g. Yoshihara & Oya 2021, Briefer et al. 2022). This creates a potential limitation to identity coding – one can expect that this information may in some cases be overridden by other, more urgent information, in cases where different types of information are not segregated in different, at least partially independent parameters (“segregation of information”; Marler 1960). In particular, some parameters of vertebrate vocalisations tend to be reliable indicators of the caller’s affective state (Briefer 2020). For instance, situations of a negative valence and/or high arousal often result in longer duration and higher frequencies, and an increased entropy and presence of non-linear phenomena in the uttered vocalisations (Briefer 2020, Gavojdian et al. 2024), which in some cases might lead to reduced vocal individuality (e.g. Rendall et al. 1998, Green et al. 2019, Osiecka et al. 2024).

In addition to the above-mentioned factors that can blur individuality, vocalisation have been shown to vary in how individualised there are depending on their function and related constraints (“social function hypothesis”; Snowdon et al. 1997). For instance, vocalisations that are aimed at specific individuals within the group, and require the sender to be identified accurately, are predicted to be more individualised than those aimed at the whole group (Lemasson & Hausberger 2011). Similarly, vocalisations emitted from close up may be less individualised than those used to communicate at distances where visual cues are absent (Xie et al. 2024). Typically, this results in affiliative calls (e.g. contact or mating calls) - often aimed at specific individuals - showing higher individuality, while agonistic and particularly alarm calls aimed at the whole group are more stereotyped (Lemasson & Hausberger 2011; Linn et al. 2021). Low individuality in calls indicating higher urgency might, at the same time, benefit the sender by inducing confusion and increasing chances that any receivers in the surrounding might come and help, as it is the case in offspring distress calls (Lingle et al. 2012). A pattern hence seems to emerge where more emotionally positive vocalisations – often produced in one-to-one affiliative contexts – could be more individualised, while those produced during negative contexts could show less individuality. In addition, this pattern might not be limited to differences between acoustically distinct call categories, but could also occur within specific call types (e.g. contact calls), which can be produced in both positive and negative contexts (such as horse whinnies, Briefer et al 2015, or pig grunts, Briefer et al. 2019).

Most ungulate species, including these commonly bred as livestock, naturally live in socially complex and often cohesive groups (Szemán et al. 2021), and exhibit prosocial behaviours such as e.g. allogrooming (Hodgson et al. 2024). Not surprisingly, they are often vocally active animals, coding (e.g. Favaro et al. 2014, Briefer et al. 2015, Maigrot et al. 2019, Briefer et al. 2022, Laliotis et al. 2023), and perceiving (e.g. Lingle & Riede 2014, Briefer et al. 2017, Baciadonna et al. 2019) important social information in their calls. Here, we tested if individuality is maintained in negative compared to positive contact calls emitted under relatively similar emotional arousal by seven ungulate species: cattle (*Bos taurus*), goats (*Capra hircus*), sheep (*Ovis aries*), domestic pigs (*Sus scrofa domesticus*), wild boars (*Sus scrofa*), horses (*Equus caballus*), and Przewalski’s horses (*Equus ferus przewalskii*). All these species show some similarities in vocal expression of emotions (Lefèvre et al. in rev), and most of them have been shown to produce individually specific calls (e.g. Lemasson et al. 2009, Briefer & McElligott 2011b, Padilla de la Torre et al. 2016, Syrová et al. 2017, Sèbe et al. 2018). However, except for cattle in which individuality was shown to be present in calls emitted in both negative and positive valence (Green et al. 2019), we do not know yet know how these two types of information interfere, and if individuality is favoured under a specific emotional valence across species.

This study explores (1) the individual information content across affective contexts, and (2) the acoustic parameters responsible for identity coding across these contexts. We predicted, based on the social function hypothesis, that contact calls emitted in negative contexts would should lower individuality due to their often more chaotic structure and their function in addressing less specific individuals.

## Methods

### Dataset

This study revisits the data from Lefèvre et al. (in rev), which integrates recordings from Padilla de la Torre et al. 2015, Briefer, Tettamanti & McElligott 2015, Briefer et al. 2015, 2019,

Maigrot et al. 2017, 2018 & Lefèvre et al. in prep). This dataset consists of 3181 contact calls emitted by seven ungulate species (cows, goats, sheep, pigs, wild boars, horses and Przewalski’s horses) by known, adult individuals. Calls were recorded in known contexts assigned to a putatively negative or positive valence based on the context of production (i.e., corresponding to fitness threatening/frustrating vs beneficial/relieving situations – such as isolation vs reunion; see e.g. Paul & Mendl 2018, Kremer et al. 2020), and validated based on behavioural as well as physiological (for domestic species) indicators of emotions in previous studies (see the details on behavioural contexts and corresponding references to individual studies in Supplementary Materials, folder “Original data” from Lefèvre et al in rev), and details on recording procedures in the individual studies (Padilla de la Torre et al. 2015, Briefer, Tettamanti & McElligott 2015, Briefer et al. 2015, 2019, Maigrot et al. 2017, 2018 & Lefèvre et al. in prep). To avoid pseudo replication, each of the calls for any given individual comes from an independent sequence.

### Acoustics analysis

All analyses were performed in R environment (v. 4.1.3). All calls were automatically analysed using the *analyze* function of *soundgen* package (Anikin 2019). Only mean values of relevant parameters with no missing entries were kept for further analysis (defined in *soundgen*, listed in Table 2), as these are the measures commonly used on the bioacoustics studies of the species. Individuals with at least five recorded vocalisations, were kept for further analysis (see Table 2, and more details on the individuals and contexts in Lefèvre et al in rev, Supplementary Materials folder “Original data”).

### Principal components analysis

First, we confirmed that the data were suitable for factor analysis, using the Kaiser-Meyer-Olkin criterion (*KMO* function, *EFAtools* package; Supplementary Material 1). We then ran a principal components analysis (PCA) on all calls of each species separately (*prcomp* fuction, *stats* package; Supplementary Materials, folder “PCA results”), to create uncorrelated components for Hs calculations.

### Information content

To investigate whether the individual information content of the calls varied with the affective valence of the caller, we used Beecher’s information statistic, Hs (Beecher 1989) – a robust, standard method to calculate bits of information carried by a signal and assess the approximate number of individuals that can be theoretically distinguished based on this signal (Linhart *et al*. 2019).

All previously obtained principal components (PC1-11) were used to calculate the Hs, using the *calcHS* function (*IDmeasurer* package). Hs was calculated for positive and negative calls separately, for all individuals with at least three calls available per valence. To test whether the putative valence (fixed factor) resulted in a change in Hs (response variable), we ran a linear mixed model with species as a random factor (*lmer* function, *lme4* package). Model assumptions were checked using the *simulateResiduals* function of the *DHARMa* package, based on the fitted model. To obtain p-values and significance of the LMMs, we used parametric bootstrap methods, using the *PBmodcomp* function (*pbkrtest* package; PBtest output). Taking the random-effects structure into account, this is a more conservative and reliable measure than a raw LRT (Halekoh and Højsgaard, 2014)

### Parameters coding individuality

To understand which raw acoustic parameters carry information about the caller’s identity in different contexts, we calculated the potential of individuality coding (PIC; Robisson *et al*. 1993) for each of the 11 parameters (Table 2), using the *calcPIC* function (*IDmeasurer* package). This was done for each species, separately for the positive and negative valence. The obtained values were then input in linear mixed models (LMMs), with the PIC value of each parameter as the response variable, valence as fixed factor of interest, and species as a random factor (*lmer* function, *lme4* package). Model assumptions and p-values were computed as mentioned above (*Information content*), and Bonferroni adjustment was used to correct for multiple testing (i.e., the p-values retained significance at 0.005).

## Results

### Information content

The information content was higher in the positive than negative context across the species (PBtest p-value: 0.046). Note that, by contrast, for the domestic horses and pigs, the Hs values were similar between negative and positive contexts, but slightly higher in negative calls (Figure 1, Table 1) - in case of the pig, this did not have an impact on the number of individuals that could be distinguished based on this signal (Table 1).

**Table 1.**
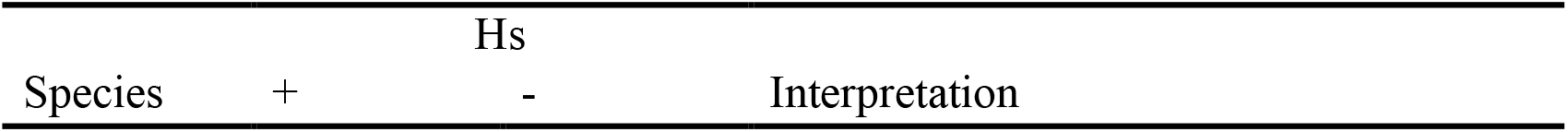
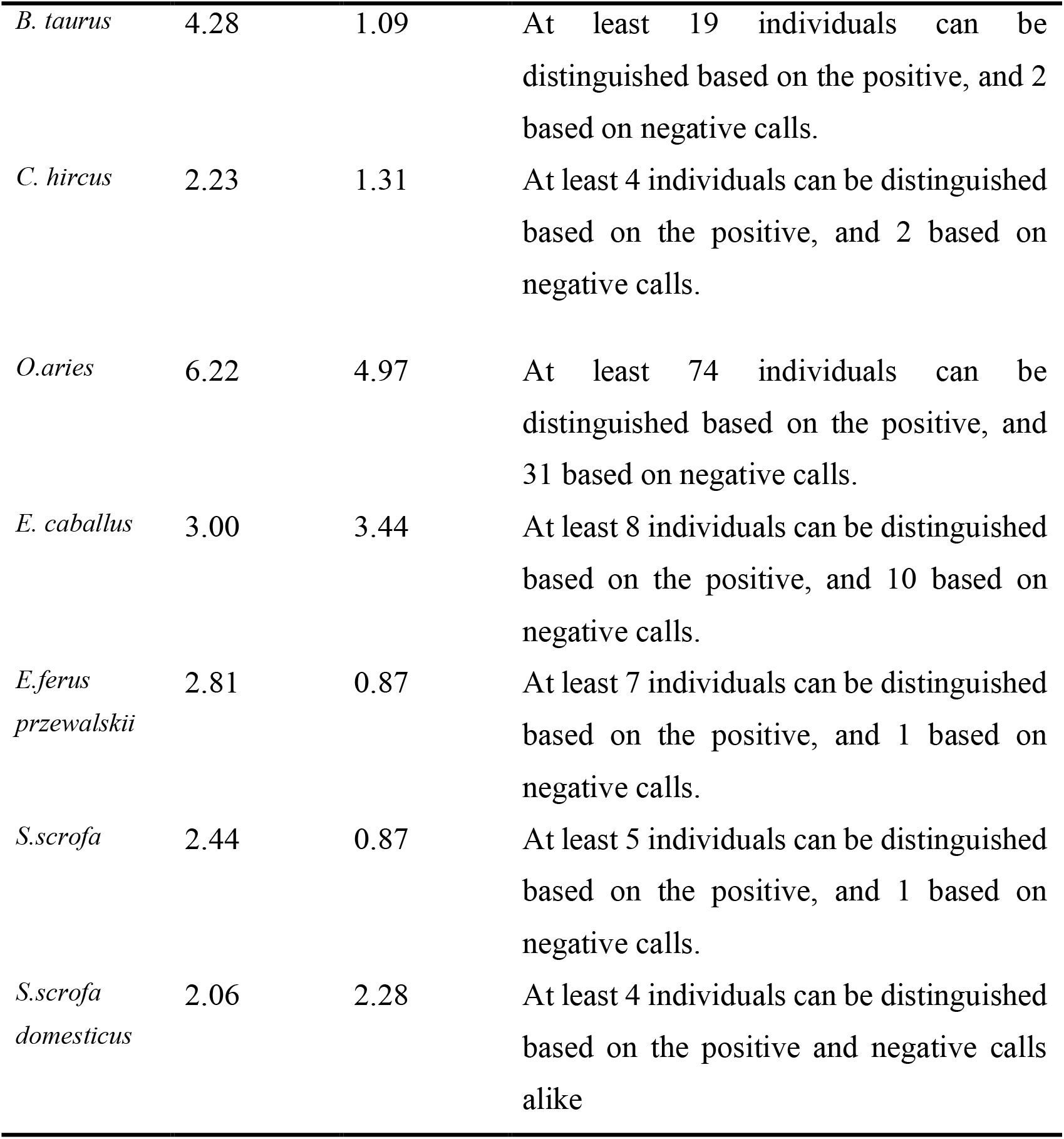
Beecher’s statistic (Hs) values for all parameters in positive and negative calls of the seven species. Approximate numbers of distinguishable individuals are calculated as 2^Hs.^

| Species | Hs |  | Interpretation |
| --- | --- | --- | --- |
|  | + | - |  |
| *B. taurus* | 4.28 | 1.09 | At least 19 individuals can be distinguished based on the positive, and 2 based on negative calls. |
| *C. hircus* | 2.23 | 1.31 | At least 4 individuals can be distinguished based on the positive, and 2 based on negative calls. |
| *O. aries* | 6.22 | 4.97 | At least 74 individuals can be distinguished based on the positive, and 31 based on negative calls. |
| *E. caballus* | 3.00 | 3.44 | At least 8 individuals can be distinguished based on the positive, and 10 based on negative calls. |
| *E. ferus przewalskii* | 2.81 | 0.87 | At least 7 individuals can be distinguished based on the positive, and 1 based on negative calls. |
| *S. scrofa* | 2.44 | 0.87 | At least 5 individuals can be distinguished based on the positive, and 1 based on negative calls. |
| *S. scrofa domesticus* | 2.06 | 2.28 | At least 4 individuals can be distinguished based on the positive and negative calls alike |

**Table 2.**
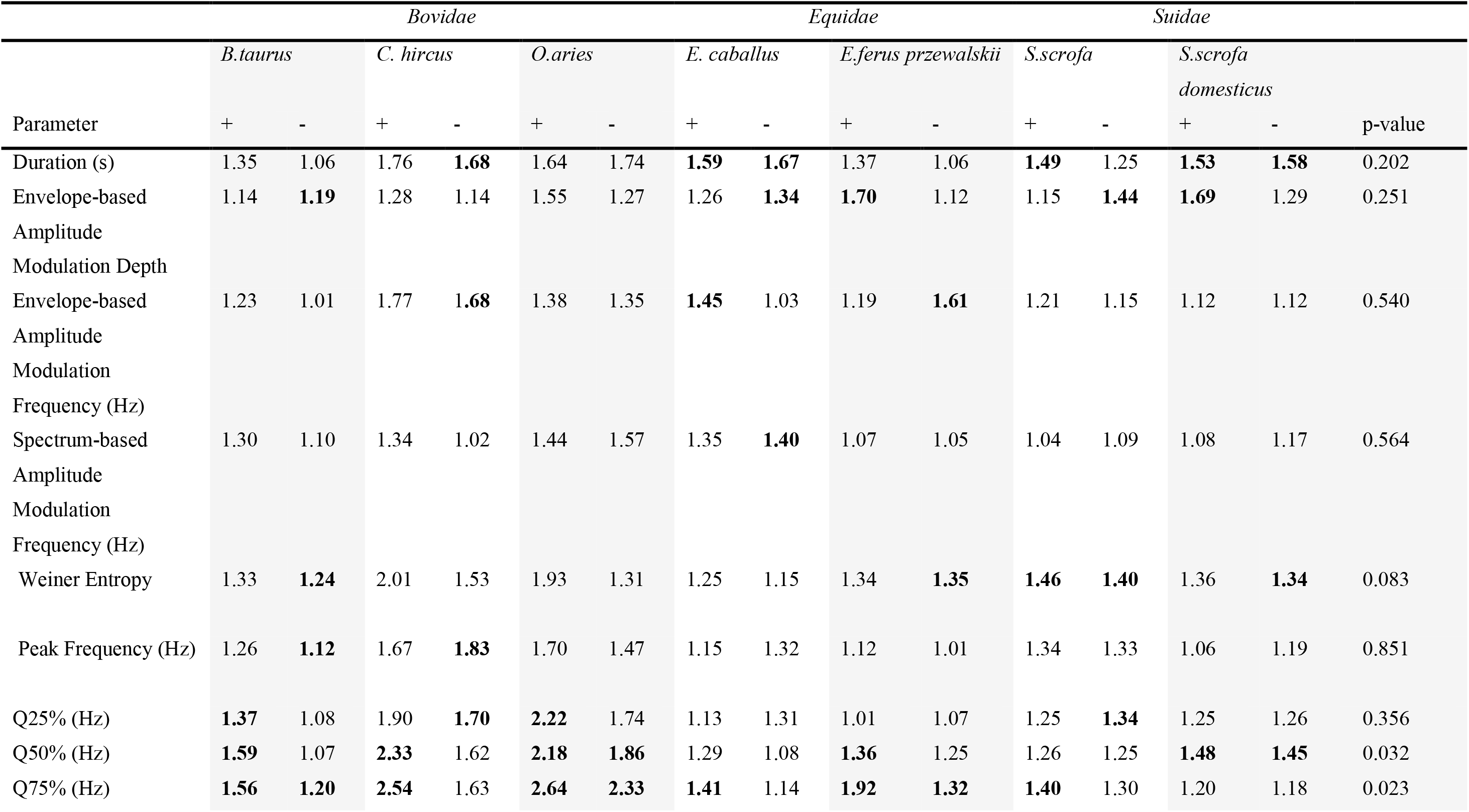
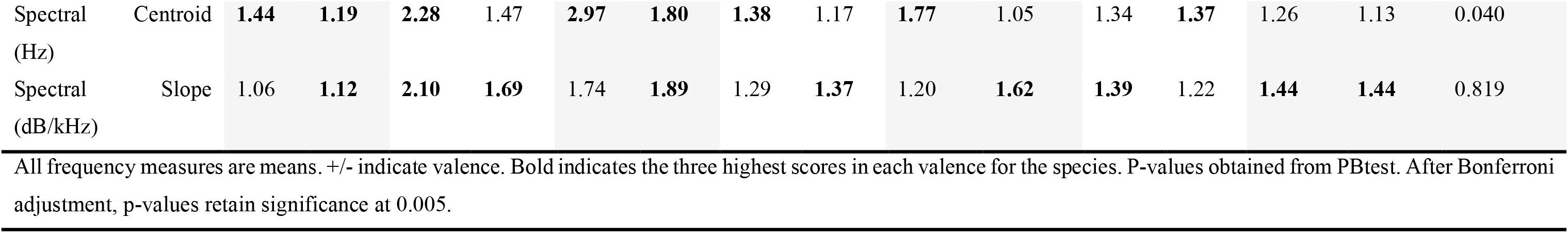
Potential for identity coding (PIC) for the 11 raw acoustic parameters. After Bonferroni adjustments (α = 0.005), PIC values showed no differences between the positive and negative contexts.

**Figure 1.**
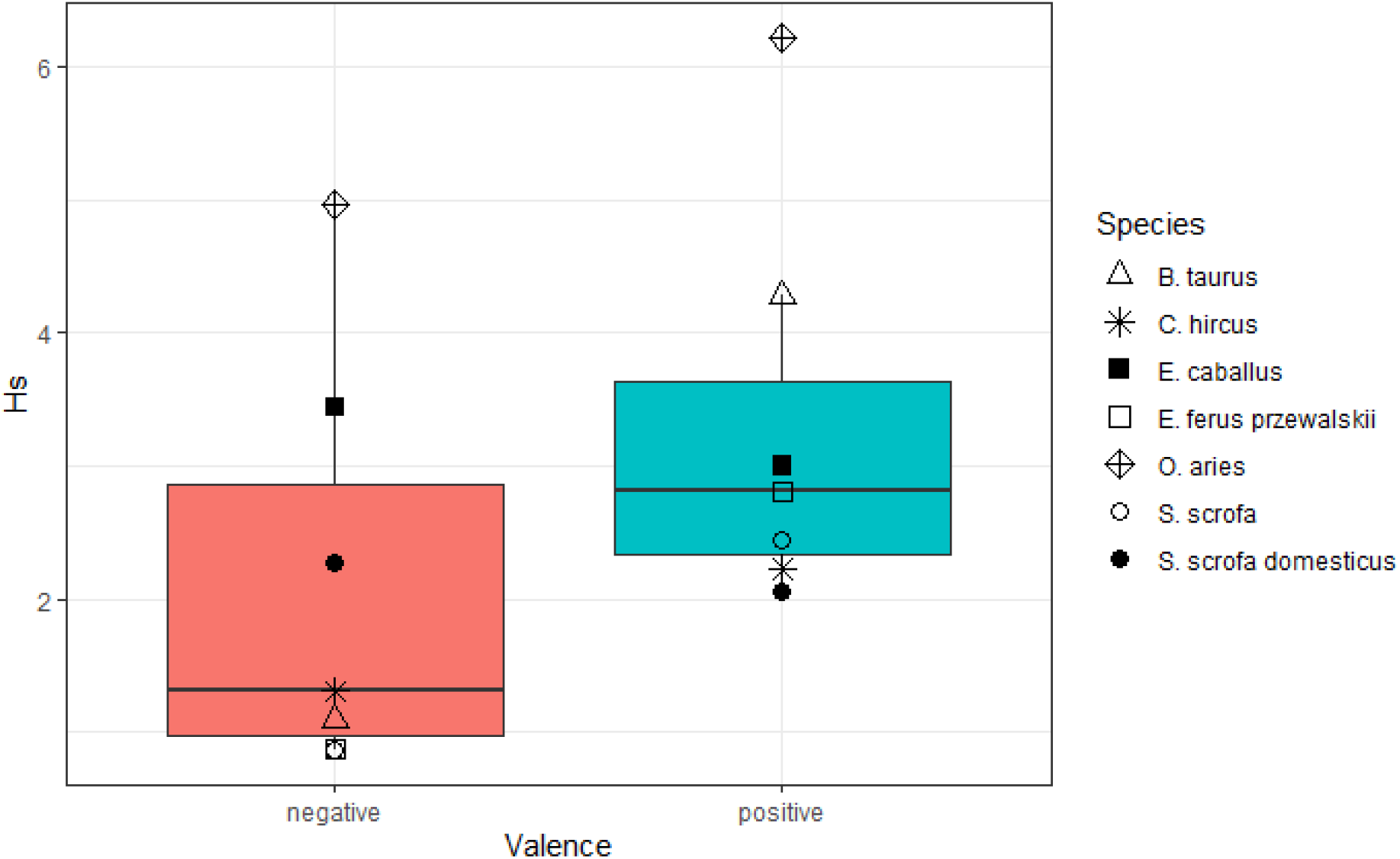
Hs as a function of valence. Boxplots represent all species pooled together, and symbols the discreet values for each of the study species. While for the domestic horse and pig the information content was slightly higher in the negative context, a general pattern seems to emerge where Hs is higher for calls of positive valence.

### Parameters coding individuality

None of the acoustic parameters showed significant differences in PIC values across positive and negative valence across the species (LMM, Table 2). For each species, at least one parameter seemed to remain a particularly stable indicator of identity between valence, such as duration spectral slope or the 75^th^ quantile of the frequency spectrum (Table 2).

## Discussion

We investigated the individuality in vocalisations uttered in putatively positive and negative contexts in seven ungulate species. Across the species, the information content about the emitter’s individuality was lower in negative than positive contexts (or remained essentially unchanged). In each species, at least one acoustic parameter seemed to have a particularly stable potential for identity coding (PIC) across the valenced states.

We predicted that the identity information might decrease in negative contexts. This prediction was based on the *arousal* and *function* of the calls. As negative vocalisations, which are often (but not always) of higher *arousal* than positive vocalisations, can be more chaotic, their information content may indeed drop (Osiecka et al. 2024), overridden e.g. by information about the level of stress experienced by the vocalising animal (Volodin et al. 2017, Corvin et al. 2024). This indeed was the case of the ungulate species studied here: in five of the seven species, the information content was lower in negative than positive calls, and in the remaining two species it remained essentially unchanged between the contexts. Vocal individuality levels may instead depend on the social *function* of calls (Snowdon et al. 1997), i.e. whether the calling animal expects the members of their social group to react (and thus needs to convey information on their identity), expects anyone to react (and thus needs to convey urgency rather than identity), or simply vocalises in an uncontrolled manner (again reflecting the urgency or arousal). Indeed, isolation calls often retain individuality (e.g. Sibiryakova et al. 2017, 2018, Green et al. 2019), while calls intended to elicit release and/or produced in situations of high arousal tend to lose this information (e.g. Keenan et al. 2020, Corvin et al. 2024, Sibiryakova et al. 2024). This does not need to mean that the individual information is lost: for example, even though the individuality decreases in negative calls of higher arousal (distress vs. isolation) produced by goitred gazelle (*Gazella subgutturosa*) and saiga antelope (*Saiga tatarica*) neonates, they still remain highly recognisable (Volodin et al. 2017). In other words, when a situation requires a reaction from a specific known individual, and thus recognition by them, it might be beneficial to keep the individual information content of the calls high, independently of the valence or arousal (Lingle et al. 2012, Wyman et al. 2022).

All calls analysed here were contact calls (also referred to as “attraction” or “isolation” calls in the literature; see Lingle et al. 2012) used to maintain social cohesion, and as such might be expected to retain strong identity information across contexts. Yet, this was not always the case. In some species (Przewalski’s horse, wild boar), the drop in information of the negative calls was so significant, that theoretically the signal emitted in a negative context did not carry enough bits of information to allow for individual recognition based solely on this signal. Calls of other species, and most notably of sheep, still allowed for recognition across multiple individuals, even when their information content dropped.

How is this individuality coded across the emotional contexts? Here, we compared the potential of individuality coding of various acoustic parameters between the contexts of opposite valence. While all species displayed at least one acoustic parameter than retained a high PIC across the contexts, the exact parameters also varied across the species, and no inter-species pattern was apparent. This highlights that species-specific patterns in communication always need to be described and considered separately.

### Practical implications

The fact that emotional contexts have an impact on vocal individuality is important for developing targeted welfare monitoring systems, e.g. precision livestock farming. Our results also indicate that, where one intends to monitor and recognise known individuals, it might be crucial to include model corrections for emotional valence in order to maintain efficient recognition. For example, acoustic capture-recapture is being considered or implemented in automated population monitoring of some species (e.g. Terry et al. 2005, Douhard et al. 2013, Longden et al. 2020), and could prove less reliable when the monitored animals are in a negative state or stressed. A baseline understanding of affective expression and its impact on vocal individuality in the monitored species would improve such population counts.

From the perspective of studying vocal individuality, correcting for affective states – or including calls emitted in various states, including positive and negative context, as well as high and low arousal – would be advisable to avoid under- or over-estimation of the commonly used individuality indices (see Linhart et al. 2019).

### Caveats and issues

The dataset is heavily imbalanced in terms of sample sizes for different species and individuals. Because of this, we had to take some imperfect decisions, such as keeping animals with only three vocalisations per context for the Hs calculations. While this might have some impact on the calculated index, it was the only way to keep Przewalski’s horses in this study.

We have only considered one dimension of emotions here, i.e. valence, but not arousal. High arousal will most likely influence vocal individuality (Volodin et al. 2017, Corvin et al. 2024). Here, we have included a mix of positive and negative situations of low to mild arousal for each species. A detailed study including both dimensions and calls of high arousal would be advisable to create a fuller picture.

It is also important to note that the real-life recognition can only be tested using playback experiments, i.e. asking the animals to distinguish between the calls of familiar vs nonfamiliar animals in different emotional contexts (e.g. Briefer et al. 2017), and likely to differ not only between the state of the caller, but also the receiver. We suggest that such experiments might help us understand the emotional impacts on individual recognition in the future.

## Conclusions

Our results indicate an impact of emotional valence on vocal individuality. This highlights the importance of correcting for emotional valence in studies and practical applications of vocal individuality.

## Acknowledgements

This work was funded by the Swiss National Science Foundation awarded to E.F.B. (Grant No. PZ00P3 148200) The authors would like to thank Andrey Anikin for his support on the soundgen package, and Anne-Laure Maigrot, Monica Padilla de la Torre and Piera Filippi for recording part of the audio material used in this work.

## Ethics statement

All acoustic recordings were collected in accordance with the current laws of the UK (goats) and Switzerland (other species), and approved by ethical committees as part of our previous studies. The experiments carried out to collect the goat recordings were reviewed by the U.K. Government Home Office inspector for Queen Mary, University of London. For the other species, experiments were approved by the Swiss Cantonal authorities (approval numbers: pigs, TG02/2014; wild boars and Przewalski’s horses, ZH011/15; sheep, ZH233/18; horses, VD2689; cattle, ZH49/2014).

## Authors’ contributions

ANO: Conceptualization, Methodology, Formal analysis, Writing – Original Draft, Writing – Review & Editing, Visualisation; RL: Conceptualization, Methodology, Investigation, Data Curation, Writing – Original Draft, Writing – Review & Editing; EFB: Conceptualization, Funding acquisition, Investigation, Methodology, Project administration, Resources, Writing – Original Draft, Writing – Review & Editing, Supervision. ANO and EFB share first authorship.

## Data availability statement

Raw data and the full code generated in this study are available at: https://osf.io/fm92j/?view_only=1297922ef1864ada87de5528d047243e

